# “*Form Follows Function*”: A New Paradigm of Pancreatic Cancer Progression

**DOI:** 10.1101/2025.08.12.667175

**Authors:** William Gasper, Giada Pontecorvi, Zhikai Chi, Sebastian Enrico, Lily Goodman, Haneen Amarneh, Evelyn Mazzarelli, Joshua Thomas, Lucia Zhang, Rachel Handloser, Seth LeCates, Ana Lucia Tomescu, Francesca Rossi, Myla Sykes, Anjali Bhatt, Nicholas Fazio, Alessandro Bifolco, Nafeesah Fatimah, Austin Seamann, Cheryl Lewis, Shelby Melton, Herbert J. Zeh, Megan B. Wachsmann, Dario Ghersi, Matteo Ligorio

## Abstract

Pancreatic ductal adenocarcinoma (PDAC) exhibits a distinctive propensity to invade nearby organs and infiltrate large blood vessels, even in the absence of distant metastasis. While the genetic and transcriptomic drivers of PDAC progression have been well studied, the mechanisms by which these molecular changes translate into functional, invasive behavior remain largely unknown.

Here, we uncover a striking level of tissue organization, characterized by previously unrecognized spatial and geometric properties within and among tumor structures. Leveraging the first large-scale, AI-assisted, human-curated PDAC atlas from hematoxylin and eosin (H&E) images, we annotated, classified, and characterized 144,474 malignant and normal structures from treatment-naïve (n=118) and neoadjuvant-treated PDAC patients (n=50). Additionally, we developed a new computational tool, *SHAPE*, to investigate PDAC aggressiveness through a comprehensive “*geometrization*” of cancer progression. Using traditional H&E-stained slides and three-dimensional (3D) tissue reconstruction experiments, we observed that invading tumor structures display an eccentric morphology with pronounced local angular coherence. These geometric and spatial properties revealed coherent architectural patterns, with invasive structures closely tracking vessels and nerves as they infiltrate surrounding tissue.

Mechanistically, integration of morphological features from 39,045 annotated tumor structures with whole-genome and RNA sequencing data revealed that PDACs with numerous eccentric structures exhibit increased copy number alterations (CNAs), loss of heterozygosity (LoH) on the p-arm of chromosome 17, and a quasi-mesenchymal/basal-like molecular subtype. Spatial transcriptomic analysis of 1,650 tumor structures from six additional PDAC patients further confirmed upregulation of invasive cellular programs within highly eccentric structures, such as epithelial-to-mesenchymal transition (EMT), angiogenesis, coagulation, and complement pathways, underscoring their infiltrative nature.

Finally, cross-validation of our AI-based method enabled a fully automated, highly interpretable computational approach to assist pathologists and clinicians in evaluating neoadjuvant chemotherapy response, predicting patient survival, and guiding chemotherapy in adjuvant settings.

Collectively, these findings deepen our understanding of PDAC progression, identify a new hallmark of tumor architecture, and pave the way for full integration of AI-driven morphology-based approaches into clinical workflows to improve the management of PDAC patients.

## INTRODUCTION

Pancreatic ductal adenocarcinoma (PDAC) remains the most lethal adult cancer, with a five-year survival rate of less than 15%^1,2^. PDAC’s propensity for early metastasis and its tendency to invade major arteries and veins locally and regionally – often precluding curative surgical resection^3^ - are key factors contributing to the high mortality rate observed in these patients^1^.

Despite significant advances in understanding the genomic alterations and transcriptomic programs that drive PDAC aggressiveness^4–7^, the mechanisms by which these molecular events manifest as invasive behavior remain unclear. Previously, we demonstrated that tumor structures act as discrete functional units with varying invasive capabilities^8^, underscoring the importance of tumor architecture in disease progression and highlighting the role of morphological properties during tissue invasion^8,9^.

In this study, we developed the first large-scale, AI-assisted/human-curated PDAC atlas from high-resolution H&E images (**Figure 1A**). Through a novel iterative human-in-the-loop approach, we generated interpretable segmentation masks of tumor and non-tumor structures (e.g., vessels, nerves, etc.). We further integrated genomic, bulk RNA-sequencing, and spatial transcriptomics data with morphological features of normal and malignant structures, unveiling previously overlooked aspects of tumor biology (**Figure 1A**).

**Figure 1.**
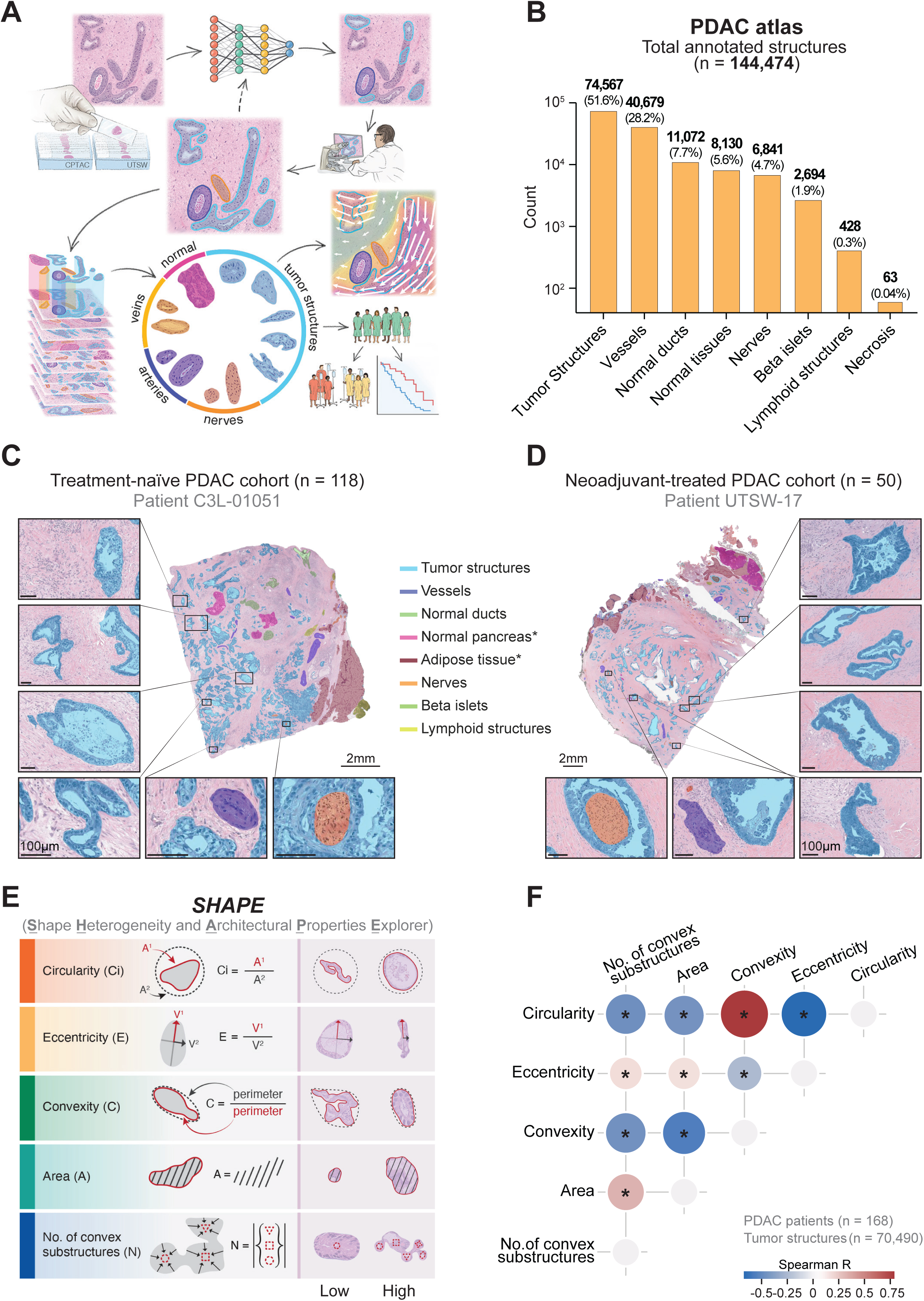
Creation of the first PDAC atlas and morphological analyses using our newly developed tool *SHAPE*. **a,** Schema of PDAC atlas creation and downstream analyses. **b**, Bar plot representing the number of annotated structures in the PDAC atlas. **c-d,** Representative images of a treatment-naïve tumor (**c**) and a neoadjuvant-treated tumor (**d**) with superimposed segmentation masks labeling different histological structures. **e**, Schema representing the five shape descriptors used by our tool *SHAPE* (<u>S</u>hape <u>H</u>eterogeneity and <u>A</u>rchitectural <u>P</u>roperties <u>E</u>xplorer) to quantify tumor structure morphology. **f**, Pairwise Spearman correlations between the five shape descriptors on 70,490 annotated tumor structures from 168 PDAC patients.

Specifically, we characterized the role of tumor architecture in PDAC progression and identified key morphological features, such as eccentricity, that significantly impact tissue invasion, vessel infiltration, and overall disease progression. Additionally, we identified a new hallmark of tumor architecture, termed “*local angular coherence”,* which combines the geometrical and spatial properties of invasive tumor structures.

Finally, leveraging our large-scale atlas, we validated a novel computational approach capable of generating interpretable segmentation masks for surgically resected PDAC specimens without human curation. As a proof of principle for its clinical applicability, we tested and confirmed its effectiveness in evaluating the pathological response of neoadjuvant therapy, assessing patient survival, and informing chemotherapy strategies in post-operative settings (**Figure 1A**).

## RESULTS

### Developing the first comprehensive PDAC atlas

To investigate how tumor architecture reflects tumor biology and impacts patient outcomes, we built the first large-scale, systematically annotated PDAC atlas using a human-in-the-loop machine-learning approach applied to standard H&E-stained slides (**Figure 1A**). This atlas consists of 173 fully annotated slides from 118 treatment-naïve patients (the publicly available CPTAC-PDA dataset^10^) and 50 fully annotated slides from 50 neoadjuvant-treated PDAC patients (UTSW dataset). Tumors from treated patients were included to evaluate the effect of neoadjuvant chemotherapy on tumor architecture, given its recent emerging role in clinical settings.

This comprehensive atlas contains readily interpretable segmentation masks, which identify normal and pathological structures, as well as regions within and surrounding the tumor area (**Figure 1B - 1D**). The ontological and spatial accuracy of AI-generated masks was ensured by two gastrointestinal pathologists (see Material and Methods) through an iterative process involving segmentation mask generation, followed by human validation and correction (i.e., human-in-the-loop). Using this approach, we annotated and meticulously curated 144,474 histological structures, including 74,567 tumor structures, 40,679 vessels, 6,841 nerves, 11,072 normal ducts, 2694 beta islets, 428 lymphoid structures, and 8,193 normal and pathological areas, including 4,828 adipose tissues, 3,028 normal pancreatic regions, 274 duodenal tissues, and 63 necrotic areas (**Figure 1B - 1D**). Tumor stroma areas were identified as the remaining tissue after this exhaustive human-in-the-loop annotation process.

### Our new tool, SHAPE, captures morphological heterogeneity in PDAC

An initial visual assessment revealed that human PDACs are composed of many approximately circular or elliptical tumor units of varying sizes, interspersed with larger tumor aggregates displaying more irregular and complex shapes.

To effectively capture this morphological heterogeneity and gain insights into PDAC progression, we developed *SHAPE*: <u>S</u>hape <u>H</u>eterogeneity and <u>A</u>rchitectural <u>P</u>roperties <u>E</u>xplorer (see Materials and Methods). This newly implemented computational tool comprises five morphological descriptors to determine the size, shape, and structural complexity of each tumor unit (**Figure 1E**). Along with area, circularity, and eccentricity, *SHAPE* includes the metric convexity. Convexity, calculated as the perimeter of the structure’s convex hull (i.e., the smallest polygon that completely contains the structure) divided by the actual perimeter, serves as a measure of structural complexity (**Figure 1E**). To further characterize complex structures, we developed a method to decompose irregular tumor aggregates into simpler subunits. This approach employs an iterative algorithm (see Materials and Methods) to identify the number of substructures (N) within each unit, which is particularly useful for analyzing these irregular tumor aggregates (**Figures 1E**).

Using these five morphological descriptors, we performed downstream analyses on 70,490 tumor structures (structures at tissue boundaries were excluded to avoid artifacts in downstream morphological analysis; see Materials and Methods) across 168 patients (treatment-naïve=118 and neoadjuvant-treated=50), finding that most tumor units are relatively small and exhibit a simple, highly convex structure. Furthermore, we noticed that morphological complexity increases with size, as indicated by a positive correlation between area and N, along with negative correlations between area and circularity, and area and convexity (**Figure 1F**). These results depict a landscape in which numerous relatively small and simple structures are dispersed among large and irregular tumor aggregates, quantitatively confirming our visual assessment.

### Identifying the key morphological building blocks of PDAC

To disentangle inherent dependencies among our five descriptors and identify archetypal structures that define PDAC architecture, we performed dimensionality reduction using principal component analysis (PCA). These analyses showed that small, simple structures are located opposite to large, irregular tumor aggregates in the PCA plot, i.e., bottom/left part vs. upper-right parts (**Figure 2A, left panel**). Additionally, PCA analysis revealed underlying morphological gradients – *circular-to-eccentric* and *simple-to-complex* - demonstrating a progression from highly circular, convex structures to irregular, complex tumor aggregates (**Figures 2A**). Altogether, these results indicate that PDAC heterogeneity, represented by a continuum of morphological shapes and sizes, can be effectively captured using our five descriptors and quantized into key morphological building blocks - discrete tumor units with distinct morphological features (**Figures 2A**). These findings led us to hypothesize the existence of distinctive architectural patterns representing combinations of structures with varying morphological characteristics (see Materials and Methods). To test this hypothesis, we applied a tensor decomposition algorithm (nonnegative Tucker decomposition; see Materials and Methods) to 39,045 annotated tumor structures from 118 treatment-naïve patients and unbiasedly identified five distinct morphological patterns (**Figure 2B**). Pattern 1, although restricted to a limited PCA region, is characterized by small tumor structures with highly circular morphology (**Figure 2B, right panels**). In contrast, Patterns 2-5 occupy broader areas of the PCA space (**Figure 2B, right panels**), encompassing a wide range of morphological features: large, irregular tumor aggregates (Pattern 2), small, convex tumor units (Pattern 3), or highly eccentric structures (Pattern 4), or large, convex tumor structures (Pattern 5).

**Figure 2.**
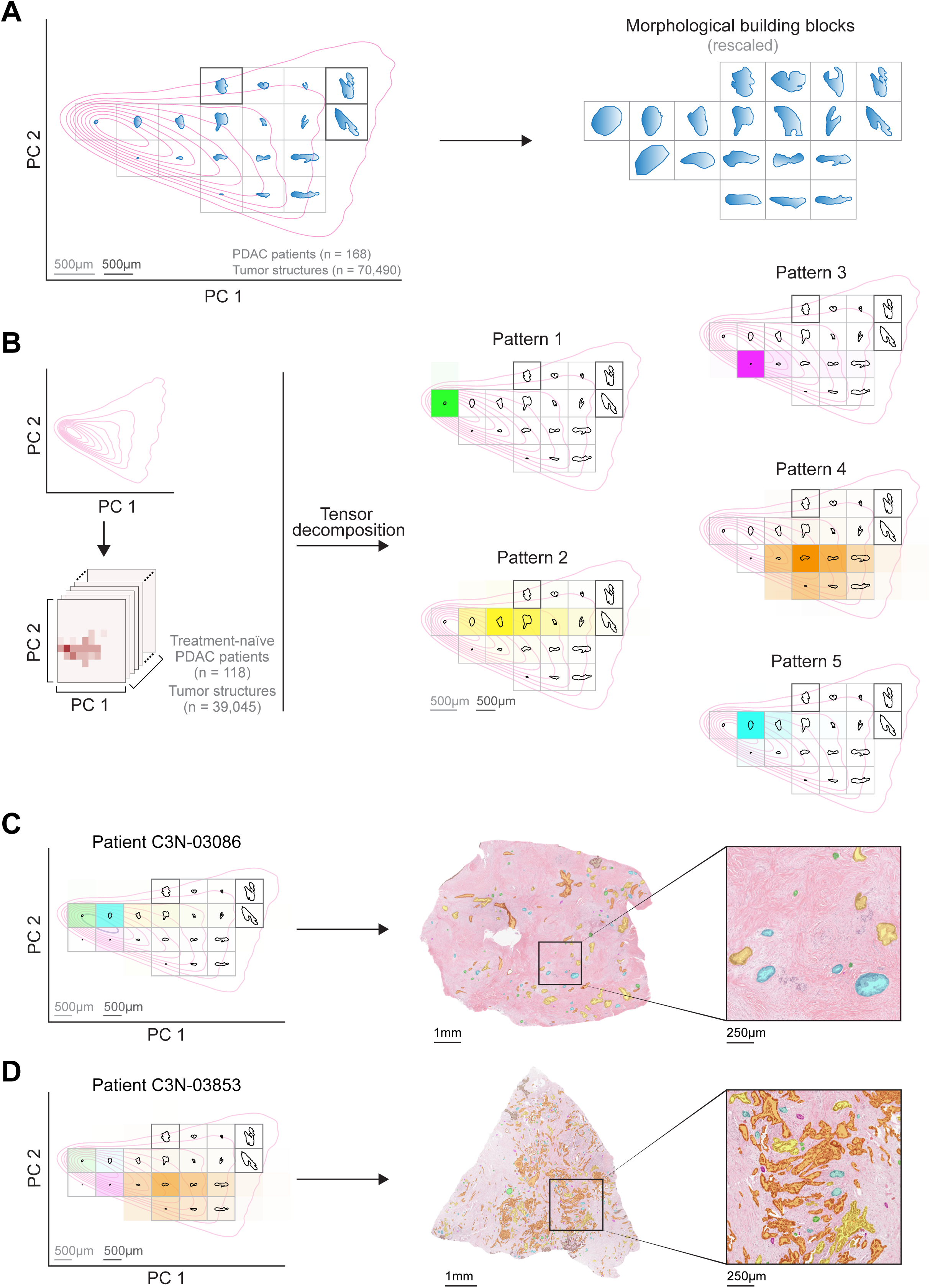
Identifying morphological building blocks and capturing PDAC shape heterogeneity using morphological patterns. **a**, Kernel density plot of PCA scores from 70,490 annotated tumor structures from treatment-naïve (n=118) and neoadjuvant-treated (50) PDAC patients. Representative tumor structures (blue) from the binned PC space (gray squares) are shown in their original size (left panel) and rescaled (right panel). **b**, Decomposing PDAC shape heterogeneity of 39,045 annotated tumor structures from 118 treatment-naïve PDAC patients into distinct morphological patterns. Creation of a tensor describing the relative frequencies of tumor structures in PC1 and PC2 bins for each patient (left panel) and application of the Tucker decomposition algorithm to identify distinct morphological patterns (right panel). Color intensity reflects the relative contribution of each PC1 and PC2 bin to each compositional pattern. **c-d**, Patient-level contribution of morphological patterns in PC space and in H&E-stained slides. Tumor structures are color-coded according to morphological pattern affiliation.

These results provided a quantitative framework for “geometrizing” tumor architecture and investigating PDAC aggressiveness through the lens of pattern contributions. Specifically, patients with a mix of highly circular and convex structures of varying sizes, along with large and irregular tumor aggregates, exhibited high contributions from Patterns 1, 2, and 5 (**Figure 2C**). In contrast, patients with small and convex structures, combined with highly eccentric tumor structures of varying sizes, showed greater contributions from Patterns 3 and 4 (**Figure 2D**).

In summary, these findings indicate that architectural patterns comprehensively capture the morphological heterogeneity observed in PDAC patients and may represent a valuable tool for gaining deeper insights into tumor biology.

### Eccentric tumor structures predict patient survival irrespective of stage

To assess whether the identified morphological patterns impact PDAC aggressiveness, we conducted outcome studies in the treatment-naïve PDAC patient cohort (n=118). Interestingly, these analyses showed that only Pattern 4 is associated with patient survival in both univariate and multivariate Cox regression analysis (**Figure 3A**). Specifically, we found that Pattern 4 and Stage IV were the only independent predictors, with higher Pattern 4 values being linked with poor prognosis, irrespective of clinical stage. These data further underscore the significance of tumor architecture on disease progression and prompted us to investigate the specific morphological features related to Pattern 4.

**Figure 3.**
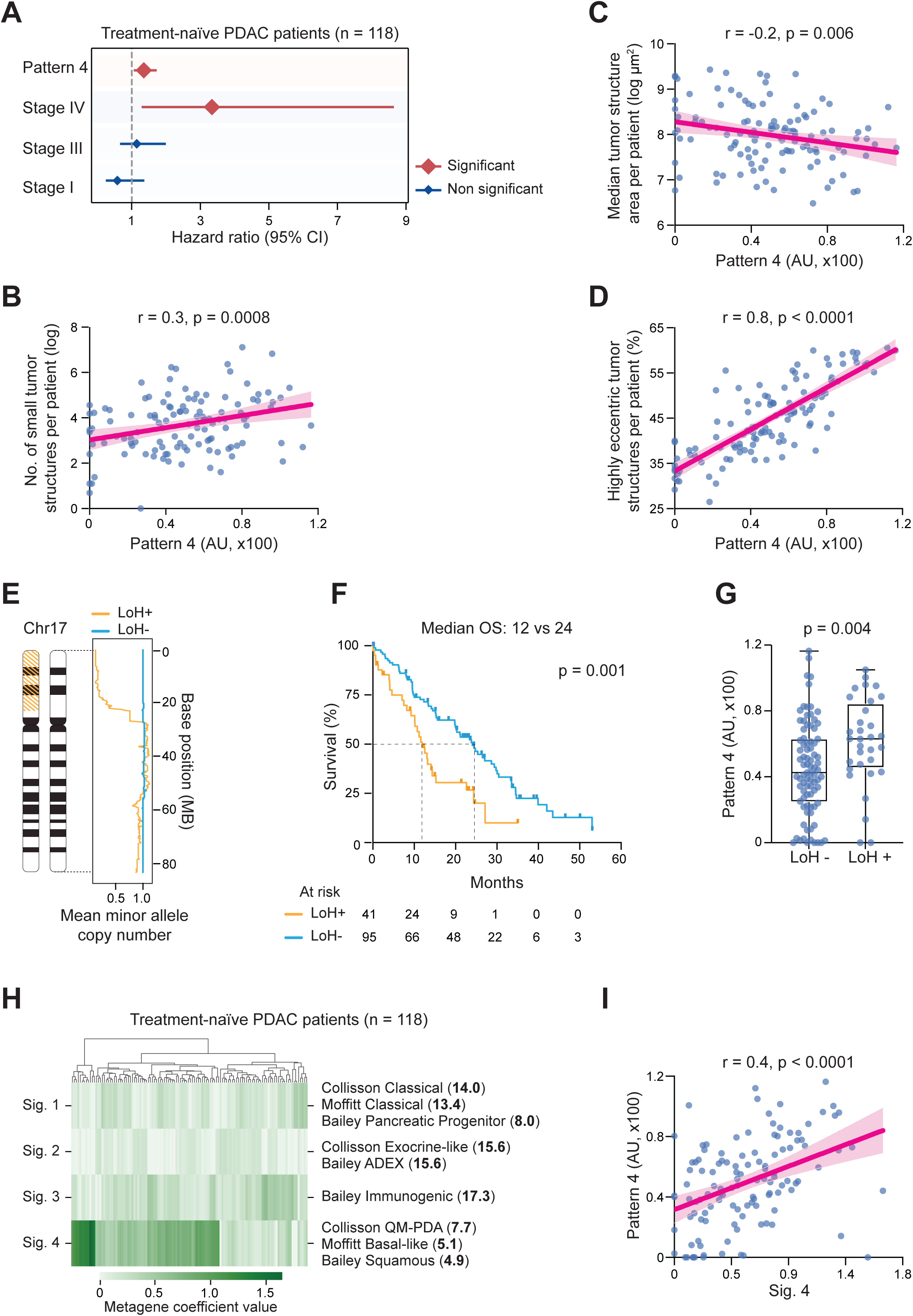
Eccentric tumor structure morphology predicts patient survival and is associated with distinct genomic and transcriptomics profiles. **a**, Plot showing hazard ratios (diamond) and 95% confidence interval (bars) from multivariate Cox regression analysis. **b**, **c**, **d**, Scatterplot visualizing the relationship between Pattern 4 values and number of small tumor structures (**b**), median area (**c**), and the relative frequency of eccentric (eccentricity > 3) tumor structures per patient (**d**). Person correlation was used to estimate the r-value and the p-value. **e**, Representation of Chromosome 17 p-arm loss (hatched in orange, left) and a plot (right) showing the mean minor allele copy number vs. base position in megabases (MB), LoH = loss of heterozygosity. **f**, Kaplan-Meier plot illustrating survival differences between patients with and without LoH for the p-arm of Chromosome 17. OS = overall survival; the p-value was obtained with the log-rank test. **g**, Box plot comparing Pattern 4 values for patients with and without LoH for the p-arm of Chromosome 17. p-value was obtained with the Mann-Whitney U test. **h**, Heatmap with hierarchical clustering showing the four gene signatures (Sig.) identified by NMF from RNA-sequencing data. Enriched PDAC molecular subtype gene sets are shown on the right. Fisher’s exact test odds ratios for gene sets are shown in bold in parentheses. **i**, Scatterplot illustrating the correlation between the Signature 4 (Sig. 4) and Pattern 4 values per patient. Person correlation was used to estimate the r-value and the p-value.

As previously shown, Pattern 4 spans a broad area within the PCA space (**Figure 2B, right panels**), comprising highly eccentric tumor structures of varying sizes, from small to large (**Figures 2B and 2D**). Shannon entropy analysis supported Pattern 4’s morphological heterogeneity, showing it has the highest entropy value across all morphological patterns (Pattern 4= 4.33 bits, Pattern 2 = 3.99 bits, Pattern 3 = 2.86 bits, Pattern 5 = 2.84 bits, Pattern 1 = 1.57 bits). Additionally, Pattern 4 displayed a positive correlation with both tumor-stroma ratios (r= 0.19, p=0.036) and with numerous highly eccentric, small tumor structures (**Figure 3B - 3D**).

Taken together, these analyses suggest that PDAC aggressiveness, regardless of stage, is associated with specific morphological landscapes characterized by a multitude of eccentric tumor structures, spanning from a few large structures to numerous small tumor units.

### PDACs with eccentric tumor structures have distinct genomic and transcriptomics profiles

To uncover the biological basis of Pattern 4 aggressiveness, we retrieved whole genome sequencing data from the CPTAC-PDA dataset and performed a series of genome-wide analyses.

As well documented in the literature^2,11^, we confirmed that *KRAS*, *TP53*, *CDKN2A*, and *SMAD4* were, in order, the most frequently mutated genes in our treatment-naïve patient population (n=118). We also observed no differences in the mutation status for TP53, CDKN2A, and SMAD4 with respect to Pattern 4 values. However, KRAS was discovered to be predominantly mutated (KRAS^G12D^, KRAS^G12V^, KRAS^G12R^, KRAS^G12C^, KRAS^G13D,^ KRAS^Q61H^, and KRAS^Q61R^) in patients with high Pattern 4 values, confirming the relatively better prognosis of patients carrying a KRAS-wildtype tumor^12^.

Additionally, given the significant role of structural variants in PDAC progression^13–15^, we investigated whether patients with high Pattern 4 values were associated with increased copy number alterations (CNAs). Thus, we retrieved CNA data and calculated the total CNA burden for each patient as the fraction of the genome that exhibited an altered copy number (see Materials and Methods). Notably, this analysis identified a positive correlation between the total CNA burden and Pattern 4 values (Pearson r = 0.24, p = 0.008), suggesting that a higher number of genomic rearrangements may drive tumor structures with aggressive behavior (i.e., eccentric structures). To further refine these analyses, we sought to identify specific genomic alterations (i.e., CNA events) associated with high Pattern 4 values. Consequently, we repurposed our computational tool, NEEP^16^, to unbiasedly identify discrete CNA events linked to poor prognosis. By applying this tool to non-overlapping 1MB regions across the genome, we found a distinctive LoH event on the p-arm of chromosome 17 (**Figure 3E**) that was linked to shortened patient survival (p=0.001, **Figure 3F**). Alterations in this region (which harbors TP53) have been extensively described in previous studies^17,18^ and were identified as the most common arm-level event in a pan-cancer analysis of TCGA data^19^. Interestingly, LoH+ patients had higher Pattern 4 values than LoH-patients (p=0.004, **Figure 3G**), further strengthening the relationship between tumor morphology (Pattern 4), genomic alterations, and clinical outcomes.

Lastly, we investigated whether Pattern 4 is associated with distinct transcriptomics profiles, given the significant contribution of different molecular subtypes to PDAC progression. We then retrieved RNA sequencing data and performed nonnegative matrix factorization (NMF), a widely used dimensionality reduction technique particularly effective for extracting gene expression signatures^4,20^. Our NMF analysis identified four signatures that fully recapitulate the three main classification systems proposed to date: Collisson^21^, Moffitt^22^, and Bailey^7^. Specifically, our Signature 1 showed enrichment for the “Classical” Collisson (odds ratio=14.0), “Classical” Moffitt (odds ratio=13.4), and “Pancreatic Progenitor” Bailey (odds ratio=8.0) signatures; Signature 2 was enriched for the “Exocrine-like” Collisson (odds ratio=15.6), and “ADEX” Bailey (odds ratio=15.6); Signature 3 for the “Immunogenic” Bailey (odds ratio=17.3); and Signature 4 for the “Quasi-Mesenchymal” Collisson (odds ratio=7.7), “Basal-like” Moffitt (odds ratio=5.1), and “Squamous” Bailey (odds ratio=4.9) signature (**Figure 3H**). These results consistently recapitulate the transcriptomics profiles of human PDACs, confirming the robustness of the NMF approach in distinguishing molecular profiles.

Next, we performed pair-wise correlation analyses to investigate whether any of the identified signatures were associated with specific morphological features. Interestingly, we identified a strong positive correlation between Pattern 4 and our Signature 4, which corresponds to the Basal-like/Quasi-Mesenchymal/Squamous molecular subtype (r=0.4, p<0.0001, Pearson; **Figure 3I**). Additionally, we observed a positive correlation between the number of small, eccentric structures – a key signature of Pattern 4 - with Signature 4 (r = 0.47, p<0.0001). These results indicate that small, eccentric tumor structures may exhibit enrichment for EMT-related genes, potentially conferring them with enhanced invasive and metastatic capabilities.

Taken together, these analyses suggest a causal link between the morphology of tumor structures (form) and their aggressiveness (function), indicating a mechanistic connection between genomic and transcriptomic alterations, PDAC architecture, and invasive behavior. As a result, these findings position tumor architecture as a novel paradigmatic framework for studying loco-regional invasion, disease progression, and patient outcomes.

### Eccentric tumor structures closely track vessels and nerves during tissue infiltration

To test the hypothesis that eccentric tumor structures express cellular programs associated with aggressive behavior, we performed spatial transcriptomic experiments (10X Visium) on an independent cohort of PDAC patients.

Under the supervision of a gastrointestinal pathologist, we selected six distinct 4.4 mm-squared regions of interest (ROIs) from four treatment-naïve patients (three different regions per patient) and four areas of similar size from two patients (two different regions per patient) who were enrolled in our rapid autopsy program (RAP). These selection criteria ensured a broad range of morphological differences while accounting for diverse clinical scenarios (early resectable vs. advanced unresectable tumors) and distinct treatment conditions (treatment-naïve vs. heavily treated patients). Images of the H&E-stained slides were acquired before any tissue processing to also assure a precise overlap between the location of each structure, its morphology, and its gene expression profile.

Using our human-in-the-loop annotation strategy, we exhaustively annotated pathological structures within H&E-stained slides and analyzed 1,650 tumor structures from these additional PDAC patients (see Material and Methods). We also reconstructed the transcriptomic profile for each structure and applied our newly developed tool, *SHAPE*, to derive their morphological properties (**Figure 1E**). Gene set enrichment analysis confirmed that the EMT pathway is enriched in small, eccentric tumor structures (**Figure 4A**), corroborating our previous findings (**Figure 3I**). Additionally, eccentric structures showed enrichment in other pathways related to an invasive phenotype, including angiogenesis, coagulation, and complement, as well as various metabolic processes, such as glycolysis, MTOR signaling, cholesterol homeostasis, and adipogenesis (**Figure 4A**). Collectively, these results support the hypothesis that eccentric structures upregulate transcriptional programs that foster infiltrative and metastatic capabilities.

**Figure 4.**
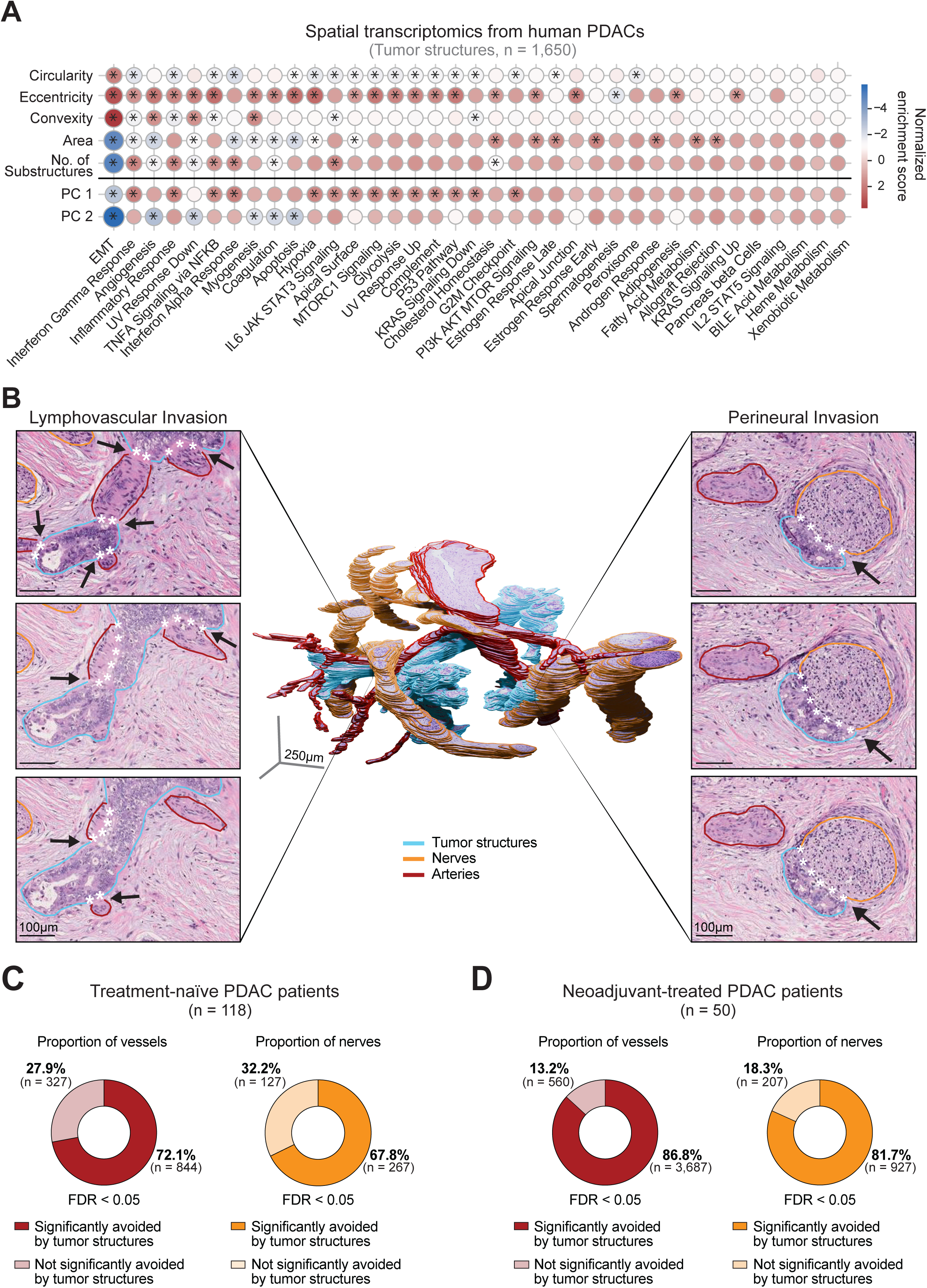
Eccentric tumor structures upregulate invasive pathways and closely track vessels and nerves during tissue infiltration. **a**, Heatmap showing gene set enrichment analysis (GSEA) results of 1,650 tumor structures from six PDAC patients for each morphological descriptor. Colors display the strength and direction of the enrichment. * indicates significant results. **b**, Serial 3D reconstruction of tumor structures (blue), arteries (red), and nerves (orange) in an invasive focus of a human PDAC (central image). Standard H&E close-ups illustrating lymphovascular and perineural invasion (white asterisks) from multiple sections. **c**, **d**, Pie charts illustrating the proportions of vessels and nerves significantly avoided by tumor structures in H&E slides from treatment-naïve (**c**) and neoadjuvant-treated (**d**) PDAC patients.

These findings prompted us to explore whether these elongated structures exhibit a preferred spatial relationship with arteries and veins during tissue infiltration. To investigate this hypothesis, we built z-stacks of consecutive 4 μm-thick H&E slides from two additional treatment-naïve PDAC patients and one RAP patient. We then examined the trajectories of small and medium-sized vessels (arteries and veins) and reconstructed the 3D trajectories of nearby tumor structures.

Surprisingly, rather than converging and immediately infiltrating vessels and nerves, tumor structures tracked these anatomical entities closely throughout the entire section (**Figure 4B, central image**). This intriguing vessel- and nerve-tracking phenomenon is in line with the high frequency of lymphovascular and perineural invasion commonly observed by pathologists when diagnosing PDAC cases (**Figure 4B, side images**). Additionally, it provides a rationale for the highly aggressive nature of Pattern 4 characterized by a high number of eccentric tumor structures - given the well-established role of lymphovascular and perineural invasion during PDAC progression, as previously reported^23–28^.

The relevance of this tracking behavior as a potential mechanism of loco-regional invasion and distant metastasis led us to analyze our treatment-naïve and neoadjuvant-treated patient cohorts (n=168), in which we had comprehensively annotated tumor structures (n=74,567), vessels (n=40,679), and nerves (n=6,841) (**Figure 1B**). By systematically examining in traditional H&E-stained slides whether eccentric tumor structures “point directly at” vessels and nerves, we further tested the vessel- and nerve-tracking capability of these invading tumor structures (**Figures 4C and 4D**). Specifically, we extended “trajectory lines” from the edges of these elongated structures, continuing their principal axis by 500 μm in each direction. We then quantified how often these trajectory lines intersected vessels or nerves and compared the observed counts to a null distribution created by randomly rotating the orientation of tumor structures 5,000 times (Materials and Methods).

Interestingly, out of 1,171 vessels and 394 nerves analyzed, 844 vessels (72.1%) and 267 (67.8%) nerves exhibited a lower intersection count than expected by chance (FDR < 0.05). This suggests that eccentric structures tend to avoid direct convergence with vessels and nerves during tissue infiltration. A similar pattern was observed in neoadjuvant-treated patients, where 86.8% of vessels (3,687 out of 4,247) and 81.7% of nerves (927 out of 1134) also showed lower intersection counts than expected by chance (FDR < 0.05). In summary, these findings corroborate the previously observed vessel- and nerve-tracking capability (3D experiment, **Figure 4B**) and reveal an unexpected yet profound spatial alignment - *local angular coherence* - among invading structures.

Taken together, these results demonstrate that highly eccentric tumor structures closely track vessels and nerves - rather than immediately infiltrating them - suggesting the existence of coordinated invasion patterns during tumor progression.

### Invasive potential: a novel architecture-based metric to assess PDAC aggressiveness

The vessel and nerve-tracking behavior of eccentric tumor structures through coordinated invasion patterns prompted us to formally test whether the angular coherence of invading tumor structures represents a novel hallmark of PDAC architecture (**Figure 5A-5B**).

**Figure 5.**
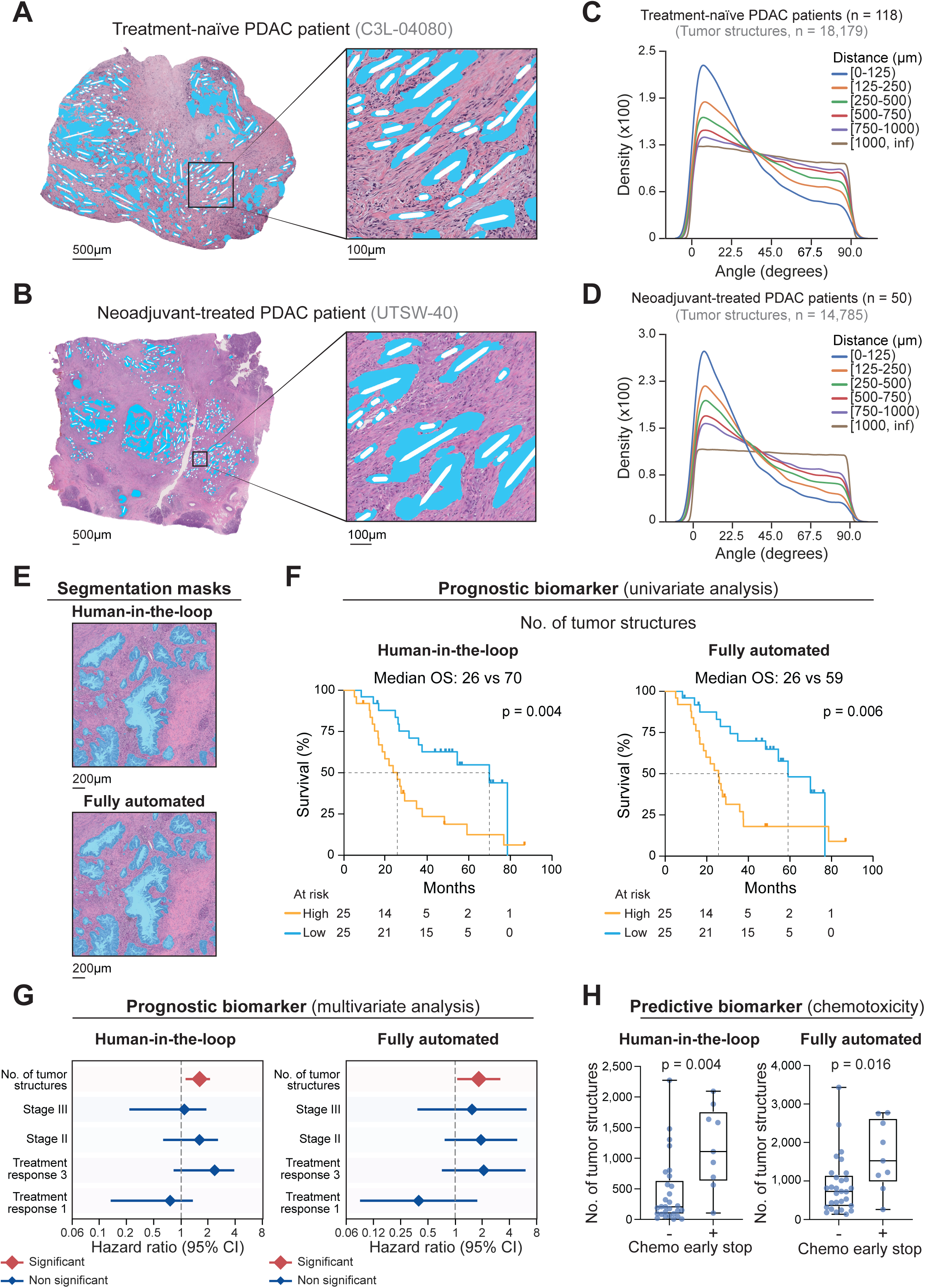
Identifying a new hallmark of PDAC architecture and leveraging architectural properties as novel prognostic and predictive biomarkers. **a**, **b**, Whole H&E slides with representative close-ups from treatment-naïve (**a**) and neoadjuvant-treated (**b**) PDAC patients illustrating the angular coherence between neighboring eccentric tumor structures. Segmentation masks (blue) show tumor structures, and bidirectional arrows (white) indicate the first principal axis for each eccentric structure. **c**, **d**, kernel density plots from treatment-naïve (**c**) and neoadjuvant-treated (**d**) PDAC patients showing the frequency distribution of the angles between tumor structure pairs at increasing distance intervals. Each interval is represented by a different color. **e**, Representative H&E close-up images comparing segmentation masks (blue) of the ground-truth, validated dataset (“Human-in-the-loop”) with the predicted masks generated by a machine learning segmentation model (“Fully automated”). **f**, Kaplan-Meier plots illustrating survival differences between patients with high and low numbers of tumor structures (binned by median) based on tumor structure counts from validated (“Human-in-the-loop”) and predicted (“Fully automated”) segmentation masks. OS = overall survival; the p-values were obtained with the log-rank test. **g**, Plots showing hazard ratios (log2 scale, diamond) and 95% confidence interval (bars) from multivariate Cox regression analysis including number (No.) of tumor structures, stage, and treatment response as covariates, based on tumor structure counts from validated (“Human-in-the-loop”) and predicted (“Fully automated”) segmentation masks. **h**, Box plots showing the number of tumor structures from validated (“Human-in-the-loop”) and predicted (“Fully automated”) segmentation masks in PDAC patients who completed adjuvant chemotherapy vs. patients who stopped early due to chemotoxicity. p-values were obtained with the Mann-Whitney U test.

To this end, we analyzed the 70,490 annotated tumor structures in both treatment-naïve (n=118) and neoadjuvant-treated patients (n=50), measuring all the angles between eccentric tumor structures (n=32,965) as a function of their distance. Intriguingly, these analyses confirmed a pronounced alignment among neighboring structures, which progressively decreases to random orientations as the distance between structures increases (**Figure 5C**-**5D**).

These remarkable findings provided the rationale for developing a new metric of invasiveness that integrates geometrical properties (eccentricity) with spatial information (angular coherence). Specifically, we combined the number of eccentric tumor structures (nES) and their median local angular coherence (mLAC) to determine the invasive potential (IP) for each PDAC case (see Materials and Methods):

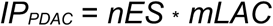

To assess the capability of this new metric to capture PDAC aggressiveness, we conducted outcome studies in our cohort of treatment-naïve (n=118) and neoadjuvant-treated (n=50) patients. These analyses revealed that IP is an independent predictor of poor survival (HR=1.7, p=0.0003, and HR=2.1, p=0.005, respectively), regardless of stage, in both patient cohorts. Notably, the multivariate Cox regression analysis that included treatment-naïve patients showed that Stage IV was no longer an independent predictor of survival (p=0.14), in contrast to when only morphological properties, such as Pattern 4, were included in the model as shown in **Figure 3A**.

In summary, these analyses indicate that this new metric, which combines spatial with geometrical properties, offers a simple, reliable method for assessing tumor aggressiveness. Furthermore, it underscores the significance of coordinated invasion patterns during tissue infiltration, establishing the local angular coherence of invading tumor structures as a novel hallmark of PDAC architecture.

### The first automated morphology-driven tool for guiding PDAC patient care in adjuvant settings

Building on our findings regarding PDAC architecture and its role in tumor progression, we sought to implement a fully automated approach capable of generating segmentation masks on standard H&E slides without requiring human curation. This AI-driven method provides high visual interpretability and is designed to support pathologists and clinicians in patient management and prognostication.

To achieve this, we leveraged our human-curated PDAC atlas, comprising over 70,490 tumor structures, and validated the first AI-based, morphology-driven approach to assist pathologists in evaluating neoadjuvant treatment responses (see Materials and Methods). Currently, tumor response scoring relies on a pathologist’s assessment of residual malignant structures^29^ - a method that, although relatively straightforward, is constrained by its visual, semi-quantitative nature. This subjectivity compromises its accuracy^30,31^, particularly when differentiating between moderate and poor responses or among varying degrees of moderate response. These limitations pose a substantial obstacle to effective treatment assessment, underscoring the urgent need for more robust methodologies to evaluate surgically resected specimens.

To address these challenges, we built a fully automated, quantitative, and highly interpretable method for determining tumor structure counts in surgically resected PDAC specimens. This AI-assisted tool generates segmentation masks that visually label tumor structures (**Figure 5E**), enabling precise quantification of the residual tumor burden following neoadjuvant chemotherapy. Validation of this fully automated approach against ‘ground truth’ data from the human-curated atlas demonstrated a strong positive correlation in predicting tumor structure counts (r=0.82, p<0.0001, Pearson). Importantly, this strong correlation translates into robust prognostic capabilities, as shown in both univariate and multivariate survival analyses (**Figures 5F** and **5G**). Notably, the AI-generated tumor structure counts emerged as the only independent predictor of prognosis, even when clinical stage and standard treatment assessments were included in the model (**Figure 5G**). Together, these data establish a compelling proof-of-principle for the utility of our AI-assisted tool in evaluating neoadjuvant chemotherapy response and in predicting patient outcomes.

To further demonstrate the clinical utility of our AI-assisted tool, we investigated whether tumor structure counts could serve as a predictive biomarker for tailoring adjuvant chemotherapy regimens. Remarkably, tumor structure count predicted early discontinuation of adjuvant chemotherapy (due to chemotoxicity), highlighting its potential to guide and optimize treatment strategies in adjuvant settings (**Figure 5H**).

In summary, these findings underscore the potential of our AI-driven tool for assisting pathologists and physicians in evaluating neoadjuvant treatment response, predicting patient survival, and informing treatment strategies.

## DISCUSSION

To the best of our knowledge, our study is the first to fully quantify PDAC morphology and “*geometrize”* tumor architecture. Collectively, our AI-driven, morphology-based approach offers new insights into the mechanisms underlying loco-regional invasion, vessel infiltration, and distant metastasis formation, uncovering previously unappreciated relationships between tumor architecture, pathology, genomic alterations, and molecular profiles.

While next-generation sequencing has transformed our understanding of tumor evolution^32^ and spatial omics have further enhanced our insights into tissue heterogeneity^9^, most studies lack a holistic interpretation of sequencing data within defined pathological landscapes. Systematic analyses that integrate architectural properties, transcriptomic profiles, and 2D and 3D spatial analyses in relation to vessels and nerves are currently missing. This gap has greatly limited our understanding of loco-regional invasion and major vessel infiltration – the two major determinants of surgical resectability. Notably, surgical resection (combined with neoadjuvant and/or adjuvant chemotherapy) remains the only curative option for PDAC patients, highlighting the critical need to elucidate the mechanisms driving tissue invasion and loco-regional vessel infiltration.

Our study aimed to bridge this knowledge gap by examining how the geometrical properties of individual structures associate with specific genomics (i.e., LoH the p-arm of chromosome 17) and transcriptomic profiles. A key innovation of our work, compared to prior studies^33–38^, is the integration of these morphological properties with global architectural features (i.e., local angular coherence), providing precise insights into how tumor structures drive tissue invasion, vessel infiltration, and overall disease progression. Specifically, our findings reveal that eccentric tumor structures in surgically resected PDAC specimens tend to track vessels and nerves during tissue infiltration. This observation aligns with previous research employing 3D reconstruction strategies to examine the microscopy behavior of cancer cells in PDAC primary tumors. Notably, studies by Noe^24^ and Kiemen^23^ demonstrated that long structures of neoplastic cells often grow parallel to normal structures such as vessels, nerves, and lobular or ductal formations. Our study builds upon this work by identifying the geometric properties of invading tumor structures, their spatial organization, associated transcriptional programs, and underlying genomic drivers. Additionally, we provide a comprehensive interpretation of PDAC architecture and tissue organization by integrating anatomical, morphological, and spatial data into a novel metric - the invasive potential - which effectively captures PDAC aggressiveness. Future research should focus on elucidating the mechanisms underlying the tropism of these invasive structures, aiming to develop therapeutic strategies that limit locoregional invasion, vessel and nerve infiltration, and ultimately, metastatic spread.

The discovery of this striking level of tissue organization in PDAC primary tumors was made possible by the development of the first large-scale PDAC atlas, comprising over 140,000 pathological structures from both treatment-naïve (n=118) and neoadjuvant-treated (n=50) patients. This AI-generated, human-curated atlas is the first to feature highly interpretable segmentation masks that accurately classify and demarcate tumor structures from normal histological entities (e.g., vessels, nerves, etc.). The creation and meticulous curation of this atlas was essential to our work, and its public availability will provide a valuable resource for the broader scientific community. We believe that making these 144,474 expert-verified annotation masks publicly available will benefit not only the biomedical field but also the machine learning community, fostering novel segmentation and classification techniques and supporting the fine-tuning of foundational image-based models, such as Meta’s Segment Anything.

The first classification of PDAC subtypes was proposed in 2011^21^, identifying three distinct molecular profiles: classical, exocrine-like, and quasi-mesenchymal. Subsequent studies have validated the existence of these PDAC subtypes and their impact on patient survival, confirming the pronounced aggressiveness of the quasi-mesenchymal/basal-like subtype^7,22^. However, molecular-based classification systems have yet to be adopted into routine clinical practice, primarily due to the lack of a straightforward pathological interpretation.

Our study also advances this area by demonstrating that histological features typical of the quasi-mesenchymal subtype, such as highly eccentric tumor structures, are easily identifiable through our morphology-based approach. This development lays the groundwork for incorporating this human-interpretable, AI-driven method into routine clinical practice. While future double-blinded prospective trials will be needed to validate the real potential of our approach in guiding patient management, the fully quantitative nature and high level of human interpretability make our method a valuable tool for assisting pathologists and physicians in clinical decision-making.

Altogether, our study provides novel insights into the biology of PDAC progression, identifies a new hallmark of tumor architecture, and introduces an innovative, highly interpretable, fully automated machine-learning approach to predict patient survival and chemotoxicity in post-operative (adjuvant) settings for PDAC patients.

## ACKNOWLEDGEMENTS

Research in the Ligorio laboratory is supported by the Cancer Prevention & Research of Texas Recruitment of First-Time Tenure-Track Faculty Members Grant (RR200023), an R37 National Institute of Health Cancer Institute Grant (5R37CA242070), American-Italian Cancer Foundation Post-Doctoral Research Fellowship to Giada Pontecorvi, PhD (2022-2024), and the Department of Surgery and the Harold C. Simmons Comprehensive Cancer Center at UTSW Medical Center.

A special thanks to the current and previous members of the Ligorio and Ghersi laboratories, in particular, Evelyn Mazzarelli, along with Rebecca Napier, Tiera Harris, and Christina Rodriguez, for their administrative support.

We acknowledge the assistance of the University of Texas Southwestern Tissue Management Shared Resource, a shared resource at the Simmons Comprehensive Cancer Center, which is supported in part by the National Cancer Institute under award number P30 CA142543.

## METHODS

### Datasets and data processing

We selected the CPTAC-PDA(*1*) cohort as our treatment-naïve group. Images were obtained from The Cancer Imaging Archive(*2*) and corresponding clinical, genomics, and transcriptomics data were obtained from the Genomic Data Commons(*3*). Whole slide images (WSIs) were included in analysis based on the ability to confidently delineate and classify histological structures. 28 low quality WSIs were excluded based on extensive tearing, occlusion, retraction artifacts, tissue hypereosinophilia, nuclear hyperchromasia, or any other phenomena that made confident cell or structure-level classification impossible. Additionally, four slides were excluded because they were entirely covered in tumor cell sheets and were not suitable for the spatial analysis presented in this work. One case was excluded based on a lack of pathologist confidence in diagnosis. One core needle biopsy was excluded due to an insufficient amount of tissue. After excluding slides, 173 slides and corresponding annotations were retained for analysis.

50 cases from the UTSW cohort were selected based on the availability of slides and detailed aggregated clinical metadata. All slides were of suitable quality and included in the analysis.

### Human-in-the-loop machine learning image annotation

One of the main objectives of our work involved elucidating morphological and architectural characteristics of PDAC, necessitating extensive annotation of WSIs. To accomplish this, we implemented a human-in-the-loop machine learning framework. Initially, six WSIs were manually annotated by pathologists (Z.C., M.W.) and trained annotators. Then, a semantic segmentation model was trained to predict tumor structures (see “Semantic segmentation of whole slide images” below). Subsequently, we applied an iterative approach where annotations were generated for a few (typically one to five) unseen slides, erroneous annotations were corrected by pathologists or trained annotators and validated by a pathologist (Z.C.) or annotation team leader (W.G.). This refined data was then added to the training pool. We utilized the CVAT image annotation platform(*4*) for annotation. Our final model predicted boundaries for tumor structures, tumor lumina, nerves, arteries, and vessels; lower frequency structures such as lymph nodes, tertiary lymphoid structures, beta cells, ductal cells, acinar cells, adipose, duodenum, and necrosis were entirely manually annotated.

We utilized the following acceptance criteria for annotations of WSIs:

- > 90% of definitive tumor pixels, defined as pixels clearly within the boundaries of a tumor structure, are within a tumor mask
- < 10% of pixels within a tumor mask are non-tumor
- > 90% of individual tumor structures belong to discrete masks (a single tumor structure cannot have multiple masks, and multiple tumor structures cannot be joined by a single mask)
- > 90% of pixels belonging to definitive normal histological structures > 8000µm^!^ are annotated
- > 90% of discrete normal histological structures belong to discrete masks

Although we utilized the above acceptance criteria, we aimed to exceed (99% correctness) these criteria on every slide. All slides in the dataset, except for those excluded due to low quality as described in “Datasets and data processing”, were annotated to meet these acceptance criteria.

After the initial manual annotation procedure, pathologist responsibility shifted to validation and training. Annotators were recruited and given the task of refining erroneous structure boundaries and received training in the form of instructional videos, documentation, and pathology workshops. Tissue structure classes and all boundary refinement work were validated by pathologists or a project manager who received extensive training from our pathologists.

### Semantic segmentation of whole slide images

We used several model architectures throughout the annotation process. Initially, we utilized a U-Net architecture(*5*). After experiencing performance plateau, we incorporated an InceptionResNetV2 encoder(*6*) in the model, and we eventually moved to a Segformer architecture(*7*). To create training data, WSIs were cropped into non-overlapping tiles and corresponding ground-truth segmentation masks were generated for each tile. During training, we incorporated various augmentation methods including stain augmentation, random rotation, random flipping, and random zooming to simulate magnification differences. For inference, WSIs were cropped into overlapping tiles, and for each tile, polygon representations of masks were obtained and stitched together to create a final representation of the whole slide. We assessed performance through tumor class intersection-over-union values.

### Shape metrics and analysis

Analyses were conducted after first converting polygon representations of histological structures into Shapely(*8*) objects. Shapely is a Python package that provides computational geometry capabilities through classes (e.g., polygons, lines, points, and others) and functions, allowing for the manipulation and analysis of spatial data.

### Circularity

Circularity quantifies the degree to which a shape resembles a perfect circle. This is accomplished by dividing the area of the shape by the area of a perfect circle with the same perimeter as the shape.

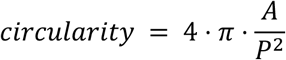

Where *A* is the area of the polygon and *P* is the length of the perimeter of the polygon.

### Eccentricity

Eccentricity was calculated as the ratio of the eigenvalues for the first two principal axes. Principal axes were defined as the first two eigenvectors resulting from principal components analysis on polygon coordinates.

### Convexity

Convexity quantifies how convex a polygon is and can be considered a measure of shape complexity. In a perfectly convex polygon, a line drawn between any two points of the polygon is guaranteed to be contained within the interior of the polygon. Convexity is quantified by comparing the perimeter of the convex hull of the polygon to the actual perimeter of the polygon, using the following equation:

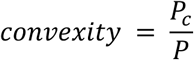

Where *P* is the length of the perimeter of the polygon and *P*_*c*_ is the length of the perimeter of the polygon’s convex hull. The convex hull of the polygon was provided by Shapely’s convex_hull attribute.

### Area

Area values were provided for polygons by the Shapely area attribute and converted to square micrometers using the WSI resolution.

### Number of simple substructures

The number of simple substructures is a measure of shape complexity and is obtained by decomposing a complex shape into highly convex substructures. To quantify the number of simple substructures, we developed a convex decomposition method inspired by and conceptually similar to Liu et al(*9*). This method involves:

1. The identification of topologically important points.
2. The identification of cut lines that can separate the previously identified topologically important points resulting in convex substructures that are more convex than the whole.

To identify topologically important points, we successively contract the polygon by 1 μm until “breaks” appear in the contractions (i.e., a contraction causes a polygon to separate into two separate polygons), and where every break indicates a topologically important point. To reduce time complexity, neighoring points are merged based on a proximity threshold (in this case, 12.5 μm). Then, edges are drawn between all topologically important points, and the edges that cross into the outside of the polygon are retained as “mutex” edges. The problem of convex decomposition then can be quite simply framed: recursively decompose the polygon by selecting cut lines scored based on cut line length (selecting the smallest cut line) and mutex weight. Mutex weights are determined by dividing the length of the shortest path between points within the polygon by the absolute distance between the points (*l_2_* / *l_1_*).

### Selection of tumor structures for morphological analysis

Tumor structures were excluded from any analyses utilizing shape metrics (circularity, eccentricity, convexity, number of convex substructures) or requiring precise determination of structure boundaries if any part of the tumor structure was within 10µm of the tissue region border. These structures were not excluded from quantification of total tumor area or total number of tumor structures.

### Stroma identification in WSIs

To identify stroma in individual WSIs, we removed all annotations from each slide’s tissue region, resulting in polygons representing stroma.

### Calculation of angle between structures

Angles between polygons with eccentricities greater than 3.0 were calculated using the following equations:

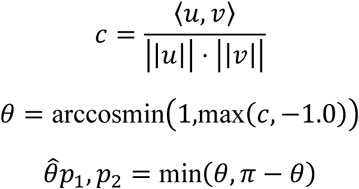

Where *u* is the first principal axis of polygon *p*_1_, and *v* is the first principal axis of polygon *p*_2_. To avoid assumptions about polygon directionality, the minimum of (*θ*, *π* − *θ*) was taken for each angle *θ*.

### Principal component analysis on shape descriptors

Shape descriptors (circularity, eccentricity, convexity, area, and number of simple substructures) were computed for 70,490 tumor structures meeting the tissue boundary criteria (previous subsection). Distributions for eccentricity, area, and the number of convex substructures were found to be right-skewed and were log-normalized and then min-max scaled prior to performing PCA. PCA was performed using the PCA class from scikit-learn(*10*).

### Identification of representative shapes for PCA bins

To provide examples of the morphological characteristics of shapes across the range of PC1 and PC2 values, we first binned the PC1 and PC2 values for the CPTAC-PDA patients into 10 equal width bins. Then, for each pair of PC1 and PC2 bins, we chose the structure closest to the centroid of the grid square bounded by the PC1 and PC2 bin edges, resulting in a representative shape for each grid square.

### Statistical analyses

Mann-Whitney U tests, Pearson correlations, Spearman correlations, and Fisher’s exact tests were calculated using SciPy(*11*).

### Data visualization

Plots were generated using seaborn(*12*), matplotlib(*13*), and GraphPad Prism. WSIs were manipulated using openslide(*14*) and Pillow(*15*).

### Nonnegative Tucker decomposition

To extract morphological composition patterns (combinations of structures with specific morphology), we utilized nonnegative Tucker decomposition to decompose joint frequency distributions describing the relative frequencies of structures, captured by their PC1 and PC2 values. To generate joint frequency distributions, PC1 and PC2 values for all CPTAC-PDA patients were partitioned into 10 equal width bins. Then, the relative frequencies of structures in each bin were computed for each patient.

After creating a tensor of joint frequency distributions with the dimensions of 118 by 10 by 10, with each 10 by 10 array corresponding to a patient, nonnegative Tucker decomposition was performed using TensorLy(*16*). We initially assessed potential core tensor rank by examining reconstruction errors in a grid search of choices for all three modes: for the patient pattern mode, ranks in the range of [1,9] were tested; for the two shape pattern modes, ranks in the range of [1,8] were tested. This resulted in only minor variation in reconstruction error for the top performing rank candidates (as assessed by reconstruction error), so we iteratively reduced the rank for each mode until redundant components in the resulting factor matrices disappeared. This resulted in a final core tensor size of 5×4×4.

### Visualization of nonnegative Tucker decomposition results

To generate PCA-space visualizations describing the tensor decomposition patterns, we utilized the resulting core tensor and factor matrices. To generate the base pattern visualizations (**Figure 2B**), which describe the occupied PC1 and PC2 spaces of patterns without respect to patient-specific values, contribution values for PC1 and PC2 were first computed by summing the products of corresponding columns in the core tensor and rows in the factor matrix (corresponding to PC1 or PC2). Then, values in the corresponding 2D PCA bins were generated by multiplying the PC1 and PC2 contribution values. To generate visualizations describing pattern activity in patients (**Figures 2C** and **2D**), the previously described base pattern values were multiplied by the normalized (divided by the maximum value of all patients for the pattern) value of the pattern for the patient.

### Survival analyses

Survival analyses were performed using the lifelines package for Python(*17*). In the CPTAC-PDA cohort we utilized stage II as the baseline variable for stage dummy variables due to a high negative correlation (Spearman’s *ρ* = −0.57, *p* = 3.55 ⋅ 10^-11^) between stages II and III. Tensor decomposition pattern values and invasive potential values were z-score normalized prior to all Cox regression analyses to make hazard ratios more interpretable. CPTAC-PDA patients with stage indicated as “unknown” were dropped from all multivariate analyses utilizing stage as a covariate.

### CPTAC-PDA transcriptomics and genomics analysis

We selected the top 3,000 highly variable genes using the highly_variable_genes function from scanpy(*18*). The number of components for nonnegative matrix factorization was chosen by examining reconstruction error for rank candidates in the range of [2,12), and then selecting the number of components corresponding to a plateau in rate of error drop.

Gene set enrichment was assessed on the top 200 genes contributing (based on the NMF H matrix value) to each component using Fisher’s exact test and significant gene sets (*p* < 0.10 after Benjamini Hochberg correction) were reported. Moffitt and Collisson gene sets were retrieved from the original publications(*19*, *20*). Bailey gene sets(*21*) were obtained by using the gene programs (provided in the supplementary data) most strongly contributing to each subtype (GP1 for pancreatic progenitor, GP2 for squamous, GP6 for immunogenic, and GP9 for ADEX), as described in(*22*).

Copy number alteration (CNA) burden was determined by taking the proportion of the genome affected by copy number alteration, using only the genomic regions for which CNA was assessed for each patient as the denominator.

To identify specific (either focal or arm-level), impactful CNA events, we first calculated the mean copy number (weighted by the proportion of the CN segment overlapping the window) for consecutive, non-overlapping 1MB windows across the entire genome. Then, we repurposed NEEP(*23*), a tool developed to identify transcriptomics features that are linked to survival, to identify CNAs correlating with patient survival in the CPTAC-PDA cohort. We observed that copy number losses for 1MB windows spanning the chromosome 17 p-arm constituted the highest impact CNAs. Based on this, we identified an arm-level loss-of-heterozygosity (LoH) of the chromosome 17 p-arm as a likely impactful, recurrent event in the CPTAC-PDA cohort. To determine chromosome 17-parm LoH status for each patient, we computed the average minor allele copy number for the 2MB-22MB region of chromosome 17, weighted by the length of the segment overlapping with the 2MB-22MB region (for example, a patient may have multiple segments of varying copy number overlapping the target region). Then, patients with a mean minor allele copy number of less than 0.5 were selected as LoH+.

### 10X Visium

To conduct 10X Visium spatial transcriptomics experiments, 4 treatment-naïve and 2 neoadjuvant-treated PDAC patients were selected under the guidance of a gastrointestinal pathologist, who reviewed H&E-stained slides. RNA extraction was performed on two 10-µm FFPE sections per sample using the RNeasy FFPE Kit (#73504, Qiagen), and RNA quality was assessed using a Tapestation system (Next Generation Sequencing Core, McDermott Center, UTSW). Samples with a DV200 score greater than 48% were selected for spatial transcriptomic analysis. According to 10x Genomics’ instructions, FFPE blocks were placed in an ice bath for 15 minutes before sectioning. 5-µm-thick tissue sections from the selected regions of interest (between 4.4×4.4mm and 6.5×6.5mm) were mounted onto the capture area of 10x Genomics Visium Spatial Gene Expression slides for FFPE (Tissue Management Shared Resource facility, UTSW). Then, slides were dried at 42°C for 3 hours, followed by overnight incubation in a desiccator at room temperature to ensure complete drying. The prepared Visium slides were then H&E stained and scanned at 20x magnification using a Zeiss Axioscan Z1 macroscope (Whole Brain Microscopy facility, UTSW). Subsequent steps to prepare sequencing-ready libraries were performed according to the manufacturer’s instructions (10x Genomics). The quality of the Visium libraries was validated using the High Sensitivity D1000 screentape (Next Generation Sequencing Core, McDermott Center, UTSW). Sequencing was conducted at Novogene using the NovaSeq platform with a PE150 strategy.

### 10X Visium data processing

Fiducial frames on WSIs were manually aligned using the Loupe Browser software (https://www.10xgenomics.com/support/software/loupe-browser) available from 10X Genomics. Images of H&E slides were annotated using our human-in-the-loop method (previously described) to determine histological structure and region boundaries. Reads were processed using Space Ranger (https://www.10xgenomics.com/support/software/space-ranger) from 10X Genomics using default options and reference resources were obtained from 10X Genomics (probe set v1.0 for GRCh38 reference genome). Raw expression counts matrices from slides were concatenated and normalized using SCTransform(*24*) as distributed with Seurat(*25*). The top 3,000 variable genes (default value) as determined by SCTransform were retained for downstream analyses.

### 10X Visium data analysis

To identify gene expression associated with morphological metrics, we first created gene expression vectors for individual tumor structures. Gene expression vectors for individual tumor structures were generated by taking the weighted average (based on proportion of the spot intersected) of expression for all spots intersecting with the tumor structure, with a minimum intersection proportion of 0.33 required for the spot to be counted, resulting in a total of 1,650 tumor structures used in this analysis. To assess relationships between gene expression and structure morphology, we performed gene set enrichment analysis (GSEAPreranked function from GSEA 4.2.3(*26*)) on lists of genes ranked by descending correlation with each SHAPE metric. For this analysis, we utilized the MSigDB Hallmarks(*27*) gene sets. We selected gene sets with an FDR q-value less than 0.25 as significant, as recommended in the GSEA documentation (https://docs.gsea-msigdb.org/#GSEA/GSEA_FAQ/).

### Creation of tumor tracking lines

To assess the orientation of elongated tumor structures (eccentricity > 3.0), we created a tracking line extending 500µm in both directions from the boundary of the tumor structure along the first principal axis.

### Assessing tumor tracking of vessels and nerves

To assess the degree to which tumor tracking lines intersected with vessels and nerves, we employed a permutation testing scheme based on random rotation of tumor tracking lines. For each target structure (individual vessels or nerves), we first recorded the number of times tumor tracking lines intersected with the structure. Then, we performed 5,000 permutations where, in each permutation, all tumor tracking lines were rotated randomly and intersection counts with the target structure were recorded. This resulted in the generation of a null distribution for each target structure, to which the actual count was compared. Then, a percentile statistic describing the percentile of the actual count on the null distribution was assessed as significant or not significant using Benjamini-Hochberg FDR correction based on the number of tests performed (number of vessels or number of nerves). Vessels or nerves that were never intersected by a tumor structure tracking line during any permutation were excluded from these analyses.

### 3D tissue reconstruction

To reconstruct the 3D structure of tumor and benign structures, we first performed serial sectioning of a PDAC tissue block (Tissue Management Shared Resource core, UTSW), and then stained each slide with H&E. H&E staining was performed on the Leica Autostainer XL Biosystem (model ST5010) based on the standard operating protocol in the Pathology and Laboratory Medicine Service at The VA North Texas Health Care System. The H&E-stained slides were then scanned using the HAMAMATSU NanoZoomer S60 Brightfield Slide scanning system (C13210-01). Then, we performed non-rigid registration of serial sections using VALIS(*28*), with default parameters. After alignment, we used our previously described semantic segmentation model to annotate tumor structures, arteries, vessels, and nerves. Annotation coordinates were used to isolate pixels belonging to structures from the surrounding tissue, giving the appearance of removal of tissue not belonging to the targeted structures (tumor, arteries, vessels, and nerves). We then utilized Blender(*29*), Python, and the Blender API to generate 3D reconstructions of each tissue block and selected areas of focus for further isolation and development. Structure annotations within foci were manually corrected using our human-in-the-loop process and final 3D reconstructions were created using Blender.

### Assessing median local angular coherence

Tumor structure trajectory vectors, defined as the first principal axis for all tumor structures with an eccentricity greater than 3.0, were calculated for all tumor structures. Then, for all pairs of structures (with eccentricity greater than 3.0) in each slide, the angle between the structures (as previously described) and the distance between the structures was recorded. To quantify this effect at the individual slide level, we computed the median angle between all pairs of eccentric tumor structures within 250µm of each other for slides with at least 10 angles meeting this criterion. After calculating the median angle between pairs of eccentric structures within 250µm for each patient, patient-level values were min-max scaled and subtracted from 1.0 to produce a final metric where increasing values represent increasing local angular coherence. Median local angular coherence was not computed for slides with less than 10 angles, and these were discarded from subsequent analyses.

### Evaluating the automated prediction of number of tumor structures

To evaluate segmentation performance on both the CPTAC-PDA and UTSW datasets, we employed 5-fold cross validation at the slide level. First, we combined and shuffled all slide identifiers for both datasets. Then, the shuffled array was split into five approximately equal sized partitions (folds). For each fold, training data was generated on the four other folds, and then the resulting weights were used to generate segmentation mask predictions on the unseen slides in the fold. As this experiment was intended to assess real-world performance at predicting the number of tumor structures, we trained models to predict only the tumor class. When generating training data, slides were tiled into non-overlapping 1024×1024 images. Then, we trained a SegFormer(*7*) architecture model with the following parameters: a batch size of 4, an initial learning rate of 6 ⋅ 10^-5^, and an Adam optimizer, for 10 epochs. Learning rate was automatically reduced on validation dataset performance plateau, and weights were retained from both the final and the best performing epoch. The best performing weights from training were then used to generate predictions on the left-out fold. For inference, WSIs were tiled into overlapping images and the resulting predictions for the slide were aggregated to produce masks corresponding to the WSI (rather than tiles).

## Code availability

Code to reproduce the results presented in this work will be released upon publication. Prior to publication, code will be made available upon reasonable request.

## Data availability

The raw data used to produce the results presented in this work will be made available upon publication.

